# Hi-C Resolution Enhancement with Genome Sequence Data

**DOI:** 10.1101/2021.10.25.465745

**Authors:** Dmitrii Kriukov, Mark Zaretckii, Igor Kozlovskii, Mikhail Zybin, Nikita Koritskiy, Mariia Bazarevich, Ekaterina Khrameeva

## Abstract

The increasing interest in chromatin conformation inside the nucleus and the availability of genome-wide experimental data make it possible to develop computational methods that can increase the quality of the data and thus overcome the limitations of high experimental costs. Here we develop a deep-learning approach for increasing Hi-C data resolution by appending additional information about genome sequence. In this approach, we utilize two different deep-learning algorithms: the image-to-image model, which enhances Hi-C resolution by itself, and the sequence-to-image model, which uses additional information about the underlying genome sequence for further resolution improvement. Both models are combined with the simple head model that provides a more accurate enhancement of initial low-resolution Hi-C data. The code is freely available in a GitHub repository: https://github.com/koritsky/DL2021_HI-C.

## I. Introduction

THE full length of DNA in a diploid human genome reaches as much as 2 m (three billion base pairs). This remarkably long chain should be condensed into 6 *μ*m cell nucleus, compacted into so-called chromatin via histone proteins further packaged into higher-order structures with distant loops and chromosome domains being formed. Beyond this complex structure, the majority of genes are preferentially located at the specific regions in the genome of a cell, e.g., the nuclear envelope, heterochromatin domains, and their activity depends on location and the corresponding epigenetic features [1]–[3]. This implies that chromatin is not a constant but rather a dynamic structure where regulation of gene expression results from properly organized genetic material in space [4]. The positioning of genes differs from cell to cell and changes during development, differentiation, and pathological processes [5].

The 3D organization of the genome has been investigated with various Chromosome Conformation Capture (e.g., Hi-C, Micro-C) techniques and high-resolution microscopy (e.g., FISH) [6]. 3C to Hi-C techniques enable the identification of statistically contact frequencies of DNA loci. Hi-C ensures the detection of loci interacting genome-widely (‘all vs. all’) to reveal the chromatin contacts in space [7]. The analysis of obtained Hi-C reads consists of their pre-processing, mapping to a reference genome, filtering, and normalization of the revealed interaction frequencies. In this way, the interaction matrix is created from raw reads indexed by row and column. This contact matrix is then processed to capture the structural entities of the genome at the kilobase-megabase scale [8].

Concordantly, the Hi-C data demonstrate that the genome is shaped into topologically associating domains (TADs) and compartments to position the genes in space properly. TADs are described as domains with a high frequency of genomic interactions. These structures ensure connections between the regulatory elements, i.e., a promoter and an enhancer; thereby, TADs contribute to transcription regulation [9]. The compartments are formed to separate active and inactive chromatin in space and connect regions through similar chromatin modifications with distal interactions. As a result, there are two types of compartments: permissive (A) and inert (B) at low resolution [10]–[12]. The TAD borders are enriched for the CCCTC-binding factor (CTCF) and cohesin binding sites [7]. The CTCF-cohesin loops ensure the close proximity of promoter and distal enhancer according to the loop extrusion model describing possible mechanisms of genome topology in 3D space [13]. Moreover, the CTCF binding sites are recently shown to participate in the TADs rearrangement after the cell division [14]. However, the formation of the A/B compartments and some TADs is not perturbed with the CTCF loss. This implies that some unrevealed mechanisms exist to organize the genome in space, and further investigations of its 3D organization are required.

The Hi-C map represents one-to-many-points and two-point interactions as stripes and insulator cohesin loops, respectively [15], [16]. Thus, Hi-C contacts reflect which distant epigenetic elements interact. Hi-C experiments capture the genome 3D structure but generate data of limited resolution, which might be insufficient to resolve contacts with high accuracy and to zoom in on interactions of small regions. The significance of these small-scale contacts can be underestimated. Thus, shedding light on them might aid in reviewing known and unraveling unexplored regulatory mechanisms.

*In silico* predictions of genome folding in space are based on the algorithms which generate contact maps from DNA sequence alone. These predictors rely on large-scale sequencing data to envision (recover) the chromatin contacts on the kilobases level, learn and train on the genomic features of the established contact frequencies for accurate results. The algorithm has to exploit the Hi-C data with high resolution to be able to recover the structures from pair contacts to loops and TADs. The solution to the issue of low-resolution Hi-C data was proposed in recent studies. The primary aim is to retrieve the high-resolution maps from low-resolution ones.

There are several computational approaches operating under this idea but adopting different principles. First, the HiC-Plus method is a neural network using a super-resolution imaging mode; second, the hicGAN method is based on Generative Adversarial Networks (GAN); third, the DeepC network utilizes both the GAN loss function and the VGG16 (trained on image data) perceptual loss function [17]–[19]. The most recent and outbreaking approach is the Variationally Encoded Hi-C Loss Enhancer (VEHiCLE), an algorithm based on variational autoencoder and adversarial training strategy equipped with four loss functions (adversarial loss, variational loss, chromosome topology-inspired insulation loss, and mean square error loss) to enhance Hi-C contact maps resolution [20].

Akita [21] is the deep convolutional neural network trained on a micro-C dataset to predict Hi-C maps of 1-megabase DNA chunks. The input DNA is processed into a binary matrix and then is transformed by a convolutional kernel with 2-kilobase bins resolution of the Hi-C map. The model is proven to accurately predict the genomic contacts from publicly available raw sequence reads of Hi-C datasets.

Here, we develop a deep learning model that can both predict chromatin contacts in the form of a Hi-C map and enhance the resolution of this Hi-C output.

## II. Methods

In this work, we aim to use information in the Hi-C contact map equipping it with information about the corresponding DNA sequence for improving the resolution of the contact map. In other words, we are addressing the following question: is it possible to develop a neural network model that is able to combine low-resolution image (or graph) data and sequence data, providing high-resolution output?

In order to do this, we propose the following pipeline (Fig. 1), which is based on using two recently developed architectures: Akita [21] which will be referred to as sequence-to-image model, and VEHiCLE, image-to-image model. The first one is a convolutional network that converts DNA sequence into a Hi-C contact map (image). The second one is a GAN-like model, which has a generator performing resolution enhancement. Firstly, we train both of them independently but with a common dataset. Secondly, we stack their outputs into a single tensor and give it to another simple “Head” model trained independently from the previous. After that, we finetune all the architecture to obtain better results. We discuss each part of the pipeline in detail below.

**Fig. 1.**
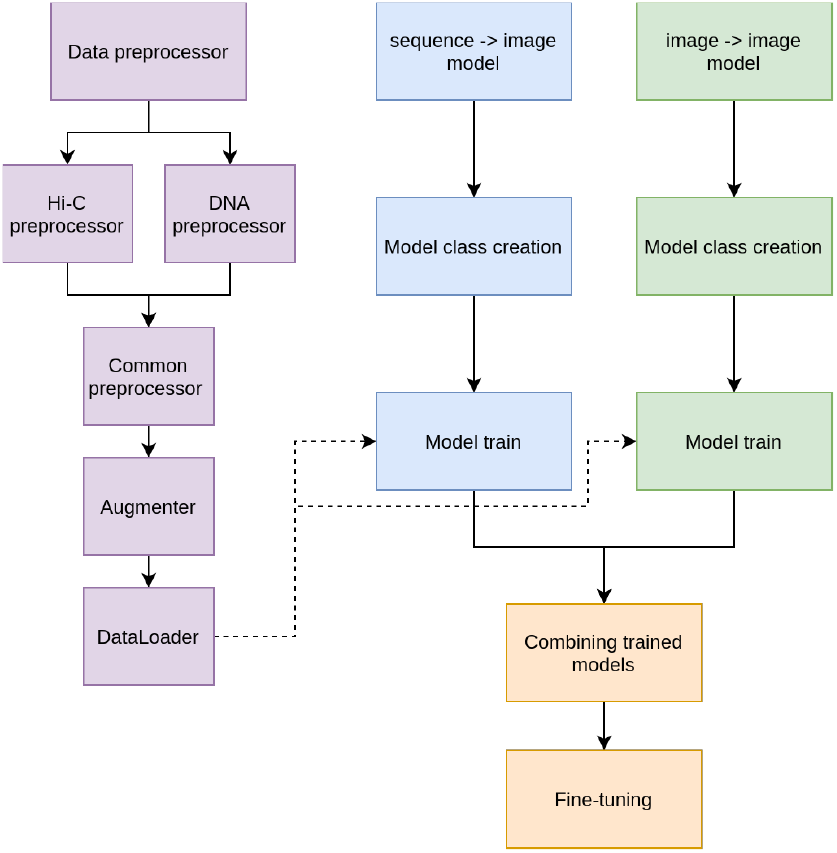
General pipeline of the project

### A. Data prepocessing

In this work, we used mouse Micro-C (hereafter, we treat Micro-C and Hi-C like synonyms) contact maps (more detailed than Hi-C) along with genome sequence data. We used GRCm38 (mm10) mouse genome assembly downloaded from [NCBI website](https://www.ncbi.nlm.nih.gov/assembly/GCF_000001635.20/), the Micro-C data was obtained from 4DNucleome database [22].

#### 1) Genome data preprocessing

The raw sequence data provided in FASTA format was split into uniform intervals of 5000 base-pairs (5kb) long by genome coordinates (in the following format: chromosome:start-end). We removed all the intervals where at least one unknown nucleotide (denoted by N) was present. Also, we removed the intervals where contact maps information was of bad quality. Additionally, we dropped the Y chromosome and mitochondrial chromosome due to poor contact data quality.

#### 2) Preprocessing of contact maps for sequence-to-image model

For this step, we fully reproduce the procedure from the Akita paper [21]. The raw contact data obtained in [Cooler](https://github.com/open2c/cooler) format was split into 200×200 pixels matrices with striding windows along the diagonal without overlapping. Therefore, each matrix corresponded to 1 million base-pairs (1Mb) genomic region; hence, each pixel reflected the contact frequency between two 5000 base-pairs loci of the genome.

Then we adaptively coarse-grain each matrix, normalize it for the distance-dependent decrease in contact frequency (diagonal-wise normalization), take a natural log, clip to (2,2), linearly interpolate missing pixels, and convolve with a small 2D Gaussian filter (sigma, 0.8 and width, 7). These steps use [cooltools](https://github.com/open2c/cooltools) library functions. The resulting contact map will be referred to as the Observed over Expected (or O/E) map in accordance with the applied normalization procedure.

Thus, we obtained 2399 examples in total, which were split into train, validation and test sets by chromosome-wise. Train set: 1912 examples from this list of chromosomes: [‘chr4’, ‘chr10’, ‘chr6’, ‘chr13’, ‘chr19’, ‘chr16’, ‘chr18’, ‘chr17’, ‘chr2’, ‘chr3’, ‘chr5’, ‘chr9’, ‘chr1’, ‘chr12’, ‘chr7’, ‘chr15’]; Validation set: 231 examples from this list of chromosomes: [‘chr11’, ‘chr14’]; Test set: 256 examples from this list of chromosomes: [‘chrX’, ‘chr8’]. Such split provides us a natural way to avoid a data leak.

Data augmentation is critical to avoid overfitting and maximize generalization accuracy on unseen sequences. Each time we processed a sequence, we stochastically shifted input sequences by up to ±11 bp, reverse complemented the DNA and flipped the contact map [21].

#### 3) Preprocessing of contact maps for image-to-image model

Since the VEHiCLE model [20] did not assume diagonal-wise normalization of contact maps, we developed another procedure of data preprocessing for the image-to-image model. The generator of the image-to-image model uses low-resolution contact maps as input and tries to enhance the resolution giving output. We created two sets of contact matrices for this purpose: (i) 200×200 pixels (target) corresponding 1Mb region as before, where each pixel reflects the contact frequency of two 5kb loci; (ii) 100×100 pixels (input) corresponding 1Mb region, but now each pixel reflects the contact frequency of two 10kb genomic loci. Furthermore, for this time, we allow matrices to be overlapped (250kb step along the diagonal or 75% overlapping) following the logic of the VEHiCLE paper where such an approach was used for correct data augmentation. Therefore, we used four times more data due to overlapping for the image-to-image model.

Then, we adaptively coarse-grain each matrix, take a natural log, normalize all values to [0..1] interval and linearly interpolate missing pixels. The resulting contact map will be referred to as Observed. We split all the data into train, validation, and test sets by chromosomes as in the previous section.

### B. Sequence-to-image model

#### 1) Original model

##### a) Architecture

Base architecture for the sequence-to-image model is taken from Akita [21]. It consists of two blocks: ‘trunk’ that takes the 1D one-hot encoded representation of DNA sequence and outputs a sequence of features, and ‘head’ transforming sequential features into 2D folding maps. The ‘trunk’ block consists of a set of 1D convolutional layers:

- Input: 1D one-hot encoded sequence;
- 1D Convolutional layer: 96 filters; kernel 11 → Batch Normalization → ReLU → Max Pooling: kernel 2;
- 1D Convolutional tower (10 layers): 1D Convolutions: 96 filters; kernel 5 Batch Normalization ReLU Max Pooling: kernel 2;
- 1D Dilated residual convolutional tower (8 layers): 1D Dilated Convolutions with increasing dilation rate + Dropout before residual connection;
- Bottleneck — 1D Convolution: 64 filters; kernel 1;
- Output: sequence of feature vectors for genomic bins.

The ‘head’ converts 1D profile into 2D map:

- 1D to 2D conversion: pairs for genomic bins from sequences are averaged using outer sum to get 2D feature map, which is concatenated with the positional encoding of the distances between bins → 2D convolution: kernel 1;
- 2D dilated residual convolution tower (6 layers): symmetrization with transposed maps is applied after each layer;
- Linear layer;
- Output: 2D maps for five datasets.

Thus, the Akita model takes as input batch of one-hot encoded sequences of length 1Mb (2^20^bp), calculates 1D profiles represented as feature vectors for 512 genomic bins of length 2,048 bp obtained from a set of 1D convolutions, which are converted into the 512×512 2D folding map through 2D convolutions.

##### b) Training

Authors trained model for 60 epochs using Stochastic Gradient Descent with momentum. The target loss function is Mean Square Error over the upper triangular portion of cropped matrices. Upper triangular indexes are taken with diagonal offset 2 and cropping size 32. They applied random shift by up to ±11 bp and reverse complement to the input DNA sequence with the following flip of the target 2D map during training as data augmentation. Next, authors used Bayesian optimization to search over the possible values of hyperparameters and chose optimizer learning rate of 0.0065, SGD momentum of 0.99575, gradient norm clipping of 10.7, and batch normalization momentum of 0.9265 as best parameters.

#### 2) Our model

##### a) Akita-like

First, we train a model with Akita-like architecture with slight changes to tune parameters for input and output data. As we have input sequences of length 1Mb and target 2D maps of size 200×200, we add an additional layer into 1D Convolutional tower block in the ‘trunk’ to get a twice higher sequence length compression. After that, ‘trunk’ output 1D profile of length 244. The resulting 244×244 map outputted from the ‘head’ block is then resized into a 200×200 tensor with bilinear interpolation. Finally, we found that the Adam optimizer with a default learning rate of 0.001 worked better in our case. We used gradient norm clipping of 10, batch normalization momentum of 0.9265, and cropping of target and predicted 2D maps by 10 on each side. We trained the model for 100 epochs with MSE loss over upper triangular values of the cropped 2D map.

##### b) Ours

Here, we replace the ‘head’ block from Akita architecture with custom graph convolutional layers. We believe this makes the model closer to the ‘physical’ representation of the chromatin as target 2D maps generally represent analogs to closeness in the 3D space. For this, we construct a new ‘head’ block. A set of Graph blocks is placed in the beginning: Graph Convolution layer → Batch Normalization layer → ReLU activation function → Linear layer → Batch Normalization. The result of the Graph block is added to the input for residual connection. Then, the resulting genomic bin features outputted from the Graph block are converted into a 2D map through the outer matrix dot product. 2D maps are then symmetrized, and batch normalization with ReLU is applied to the result. The final output is obtained after applying a single Linear layer.

For the graph convolution layer, we use a graph transformer operator described in [23]. Multi-head attention is calculated as:

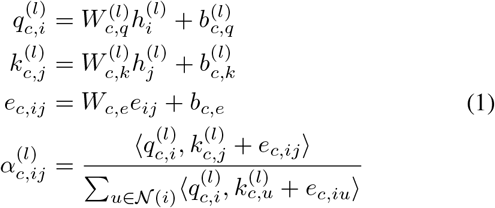

where 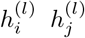 are source and distant input node features transformed into the query vector 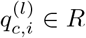 and key vector 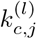, respectively, using trainable parameters 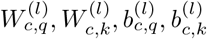 *e_ij_* are edge features further encoded through trainable *W_c,e_*, *b_c,e_*; 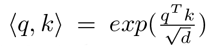 is the exponential scale dot-product; *c* is for index of the head; d is the hidden size of head. Next, message aggregation is performed through concatenating of weighted sums from each head:

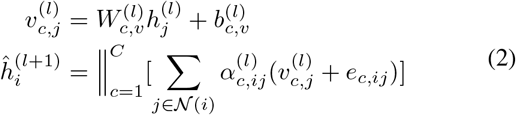

Here, we represent each node as a feature vector of the corresponding genomic bin. We use embeddings of dimensionality 8 from positional encodings |*i − j*| of the distance between genomic bins to get edge feature vectors. We use the size of the hidden dimension of each head equal to 64, the number of attention heads equal to 2, and the dropout rate of input nodes to each attention equal to 0.3.

##### C) Image-to-image model

The model VEHiCLE (short for Variationally Encoded Hi-C Loss Enhancement) from the paper [20] has the following training approach:

First, VEHiCLE incorporates a variational autoencoder that extracts biologically meaningful features from Hi-C data. Second, VEHiCLE’s decoder network is engineered to provide an easy-to-use generative model for Hi-C data generation, which smoothly maps user tunable, low dimensional vectors to Hi-C contact maps independent of any low sampled input. Third, VEHiCLE incorporates a biologically explicit loss function based on Topologically Associated Domain identification to ensure accurate downstream genomic analysis.

###### 1) The training loss

The loss that is used for training has four components: adversarial loss, MSE-loss, VAE-loss, and insulation loss. Let us explain each component.

Generative Adversarial Network consists of two components: Generator *G*, which takes samples from an input distribution and generates enhanced matrices, and Discriminator *D*, which tries to classify whether its inputs are real high-resolution Hi-C samples or enhanced resolution Hi-C samples. The adversarial loss is defined as follows (see Fig. 3 for details):

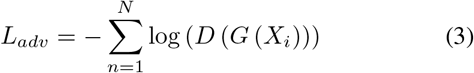

where *X_i_* are low-resolution data.

**Fig. 2.**
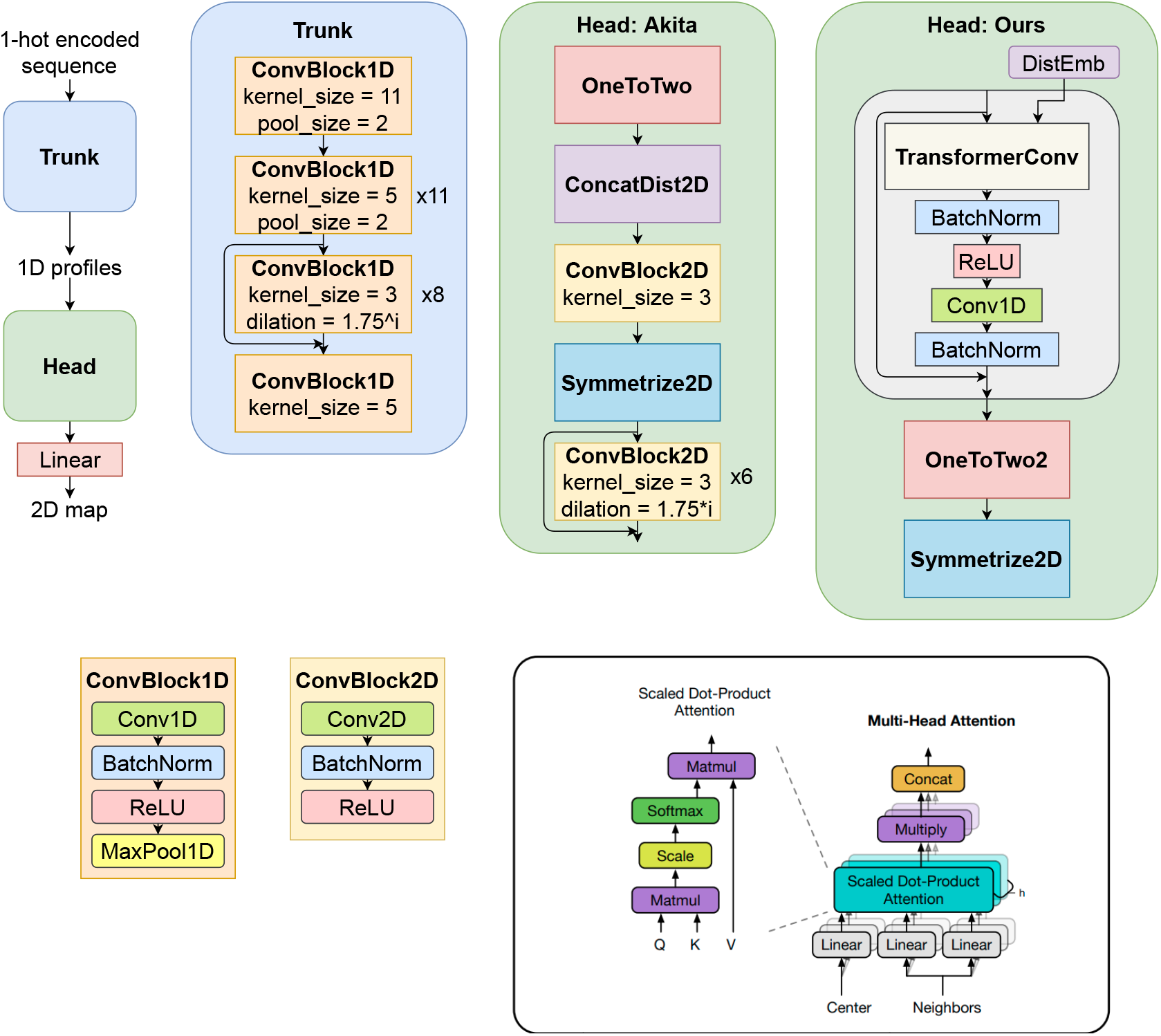
Sequence-to-image model architecture. (Image for Attention block is taken from [23].)

**Fig. 3.**
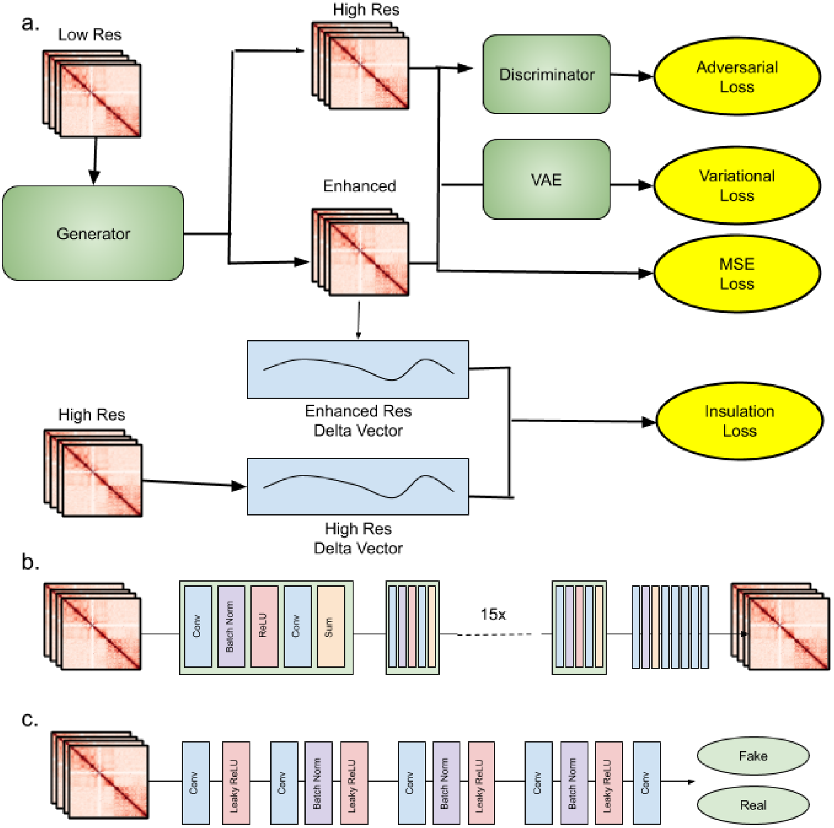
Pipeline of the image-to-image model: (a) overview of training strategy, (b) generator architecture, (c) discriminator architecture.

Bin-wise mean square error loss contributes to maintaining visual similarity between enhanced and target Hi-C contact matrices:

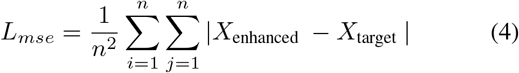

A topologically associating domain (TAD) is a self-interacting genomic region, meaning that DNA sequences within a TAD physically interact with each other more frequently than with sequences outside the TAD. Insulation loss is biologically inspired and connected to the identification of TAD boundaries using the sliding window technique and computing pseudo-derivative. During the training procedure, the insulation loss is computed with the use of a neural network *D_vec_*.

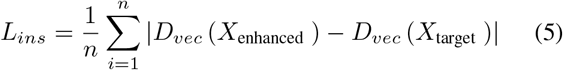

Variational loss is computed by taking the mean differences between the latent feature vectors of the enhanced Hi-C contact matrix and target high-resolution Hi-C contact matrix. These latent feature vectors are computed using variational autoencoders, which aim to condense data into lower-dimensional space, providing smooth representations.

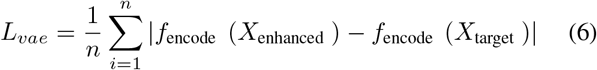

The final loss is the linear combination of the four losses: *L*_tot_ = *λ*_adv_ *L*_adv_ + *λ*_mse_ *L*_mse_ + *λ*_vae_ *L*_vae_ + *λ*_ins_ *L*_ins_ with coefficients *λ_adv_* = 0.0025*, λ_mse_* = 1*, λ_vae_* = 0.01*, λ_ins_* = 1.

###### 2) Our changes in VEHiCLE

GAN architecture was not changed since it consists of convolution blocks that do not depend on the input resolution. However, as in original implementation, the resulting image was cropped by 6 pixels on each side, so for 200×200 input image we get 188×188 as a result.

However, the VAE part has linear layers, bottleneck and transposed convolutions. To keep the latent dimension the same size — 4608 — we had to remove one convolution layer from the encoder and decoder respectively. The final architecture become 6 convolutional layers with kernel counts: [32, 64, 128, 256, 256, 512] with minor adjustments in out padding configurations.

##### D. Head model

All the aforementioned procedures can be briefly expressed with an inference scheme (Fig. 4), where the head model plays a crucial role and its architecture describing below.

**Fig. 4.**
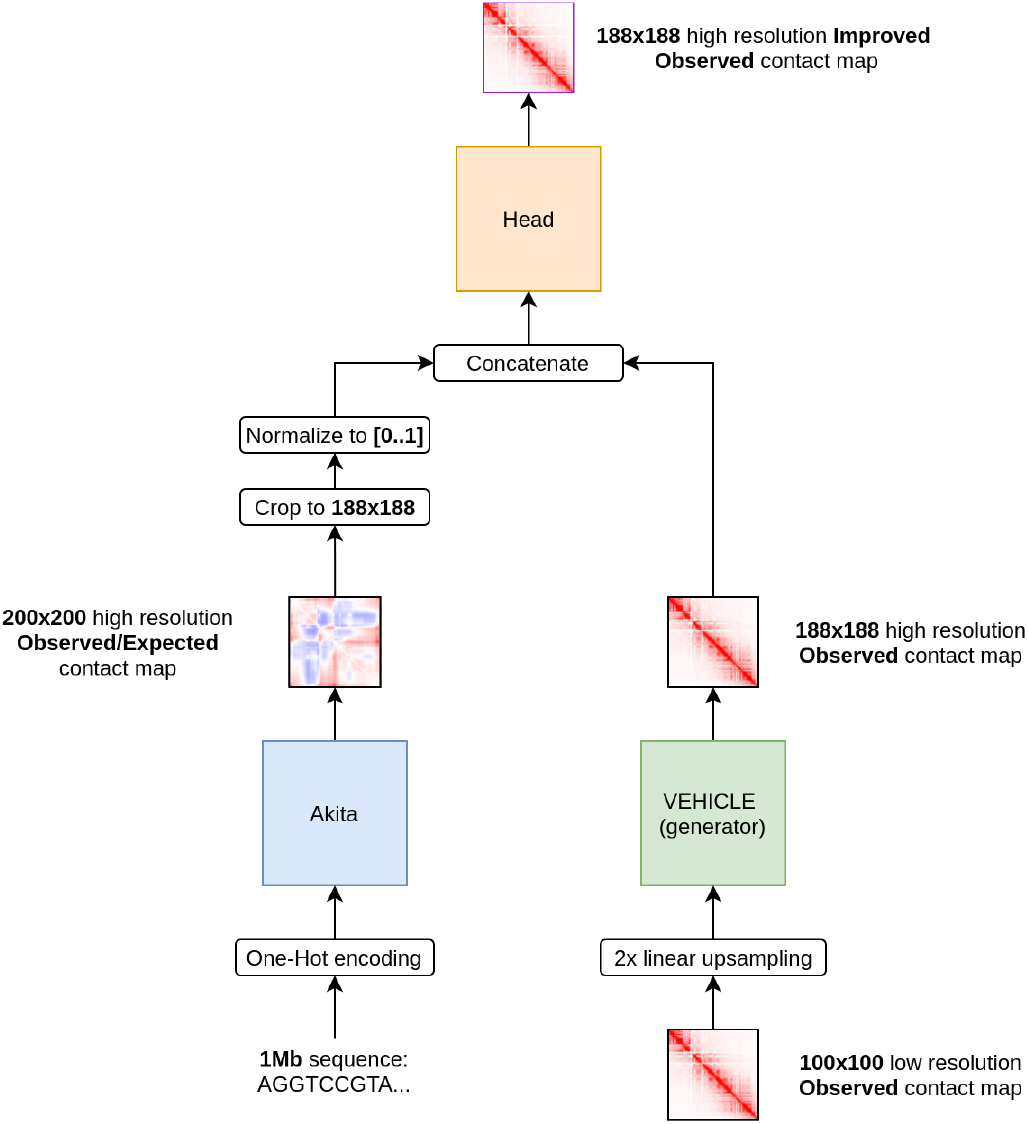
Inference process scheme

We decided the head architecture should be a convolutional net with symmetrizing procedure inside, where the symmetrization is taken from Akita model. We tried convolutional nets of different depth and kernel sizes, seeking the best one giving the highest Pearson and Spearman correlation metrics (see below). As the result we chose the architecture of 2 (3×3 kernel with padding=1) convolutional layers with kernel counts: [2→16, 16→1] with ReLU activations (what is rather natural considering that output is [0..1] normalized). After each layer the symmetrization procedure occurs.

##### E. Validation metrics

We utilize 5 reproducibility metrics (as in Vehicle paper) pulled from Hi-C resolution enhancement papers: Pearson Correlation Coefficient (PCC), Spearman Correlation Coefficient (SPC), Mean Squared Error (MSE), Mean Absolute Error (MAE) and, one specific for Hi-C task, Stratum Adjusted Correlation coefficient (SCC). For computing SCC coefficient we adopted a code from the [hicreppy](https://github.com/cmdoret/hicreppy/tree/master/hicreppy) library.

## III. Results

### 1) Sequence-to-image model

We trained two versions of Sequence-to-image models: one with an Akita-like head and the other with transformer graph convolutions with parameters described in Section Sequence-to-image model. Model performances are shown in Table I. Visual comparison of maps predicted by sequence-to-image models with Akita-like and graph-based head blocks are provided in Fig. 5.

**TABLE I.** Akita model performances

| Model | Set | MAE | MSE | Pearson | Spearman |
| --- | --- | --- | --- | --- | --- |
| Akita-like | Train | 0.220 | 0.092 | 0.778 | 0.760 |
|  | Val | 0.273 | 0.157 | 0.658 | 0.676 |
|  | Test | 0.283 | 0.151 | 0.611 | 0.585 |
| Ours | Train | 0.224 | 0.089 | 0.784 | 0.738 |
|  | Val | 0.276 | 0.158 | 0.659 | 0.647 |
|  | Test | 0.283 | 0.149 | 0.611 | 0.559 |

**Fig. 5.**
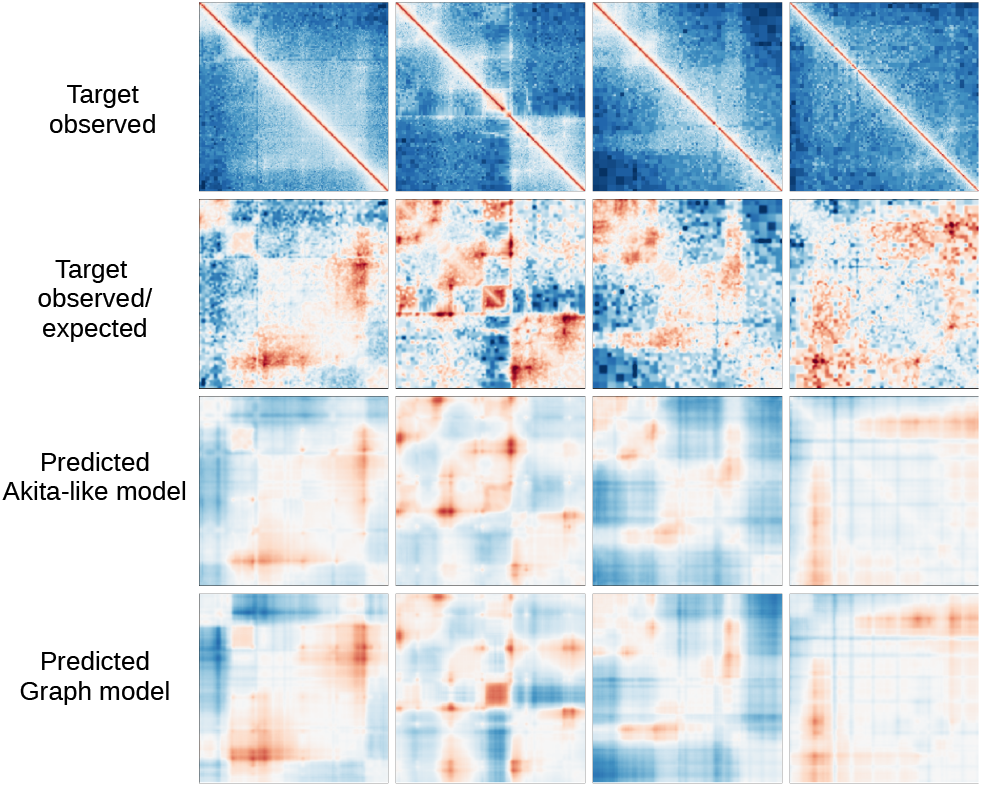
Examples of target Hi-C maps and predictions from sequence-to-image models.

### 2) Image-to-image and hybrid

The Image-to-image (VEHiCLE) part pre-trained on our data can be considered as a SOTA with which to compare the results of our model, and the hybrid model performs worse (Fig. 6). The main problem here lies in the diagonal elements that are blurred out to the thick line because the Akita model does not learn diagonals, and during the preprocessing, they are approximated with the gaussian kernel. Consequently, this step presents a potential room for improvement.

**Fig. 6.**
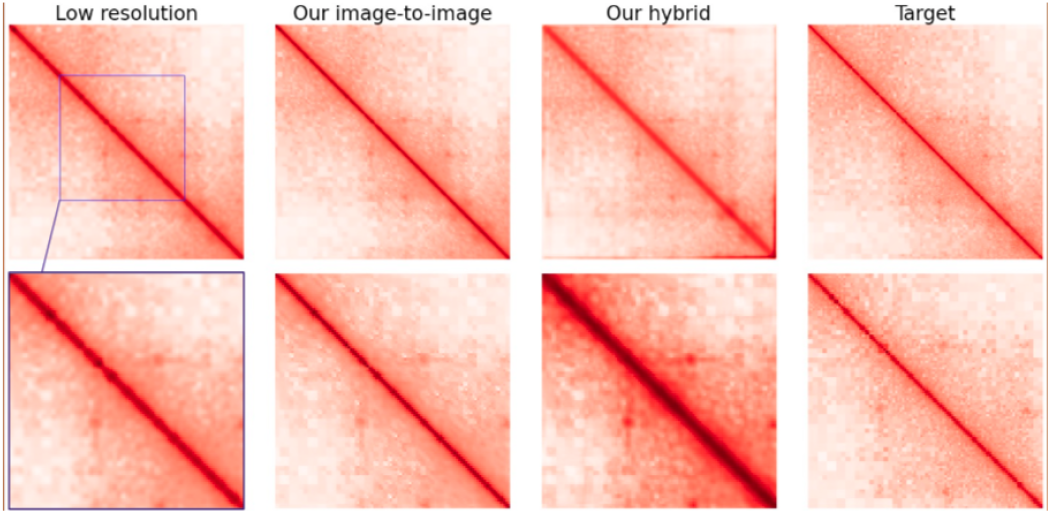
Examples of different Hi-C maps. On the top panel some region with point features is depicted. On the bottom panel the fragment corresponding to purple square is zoomed.

We implemented multiple approaches, including pre-heating the head and finetuning afterward, but did not achieve significantly better results (Fig. 8). We suggested that, after some training, the convolutional head would switch off one of the networks to get all the profit from the other, but it did not happen. To check it, we have applied zero tensors as an input for one part while the input of another part remained valid and obtained worse results than the combined (Fig. 7).

**Fig. 7.**
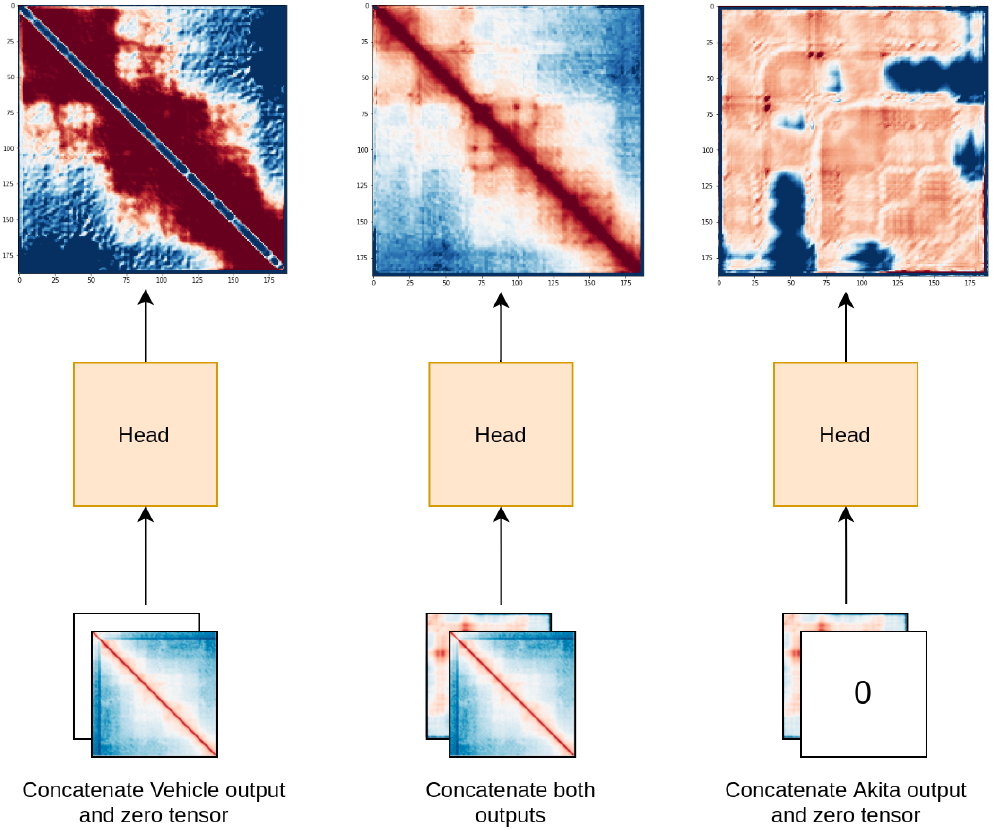
Behavior of the network head output in case of zerofying one of the backbone outputs.

**Fig. 8.**
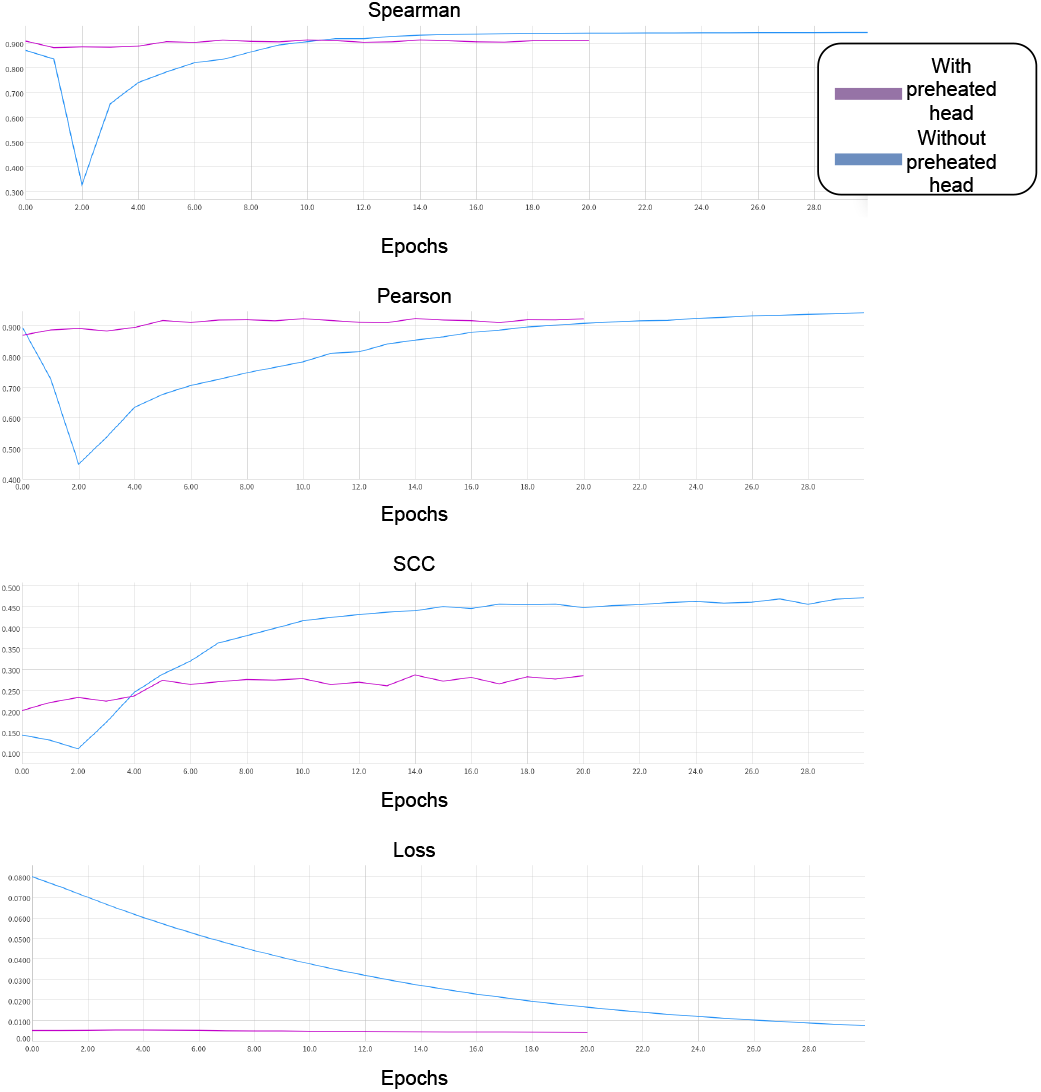
Metrics and loss on validation set for model with and without preheated head. Preheated head increases training stability at the beginning, but non preheated head shows better final results.

Thus, one can conclude that the total network learns from both backbones outputs but can not leverage from it. The reason might be in the different distributions on each output type and the small depths of the convolution head not allowing it to adjust.

To compare our results with the baseline, we used models HiCPlus, DeepHiC, HiCSR, and VEHiCLE. Since we did not have enough resources to train them on our data, we used pre-trained weights on the human genome. This dataset differs from our mice genome slightly but noticeably decreases metrics (Table II). Of note, GAN-based VEHiCLE shows much better generalizability than the previous generations of algorithms.

**TABLE II.** Metrics comparison with baselines and SOTA

| Model | MAE | MSE | Pearson | Spearman | SCC |
| --- | --- | --- | --- | --- | --- |
| Downsampled | 0.0373 | 0.0024 | 0.9491 | 0.9244 | 0.553 |
| HiCPlus | 0.1104 | 0.0189 | 0.6628 | 0.7121 | 0.1189 |
| DeepHiC | 0.0946 | 0.0178 | 0.79 | 0.6949 | 0.1801 |
| HiCSR | 0.1252 | 0.0226 | 0.7165 | 0.7155 | 0.1519 |
| VEHiCLE | 0.0807 | 0.0148 | 0.9181 | 0.9096 | 0.407 |
| img-to-img | 0.0311 | 0.0017 | 0.9551 | 0.9257 | 0.569 |
| Hybrid | 0.0327 | 0.002 | 0.9473 | 0.9247 | 0.5201 |

## IV. Discussion

### 1) Combined model

The results show that the VEHiCLE model by itself provides better results than any of our hybrid models with different head block designs and different train procedure designs.

We also explored the behavior of the head block of the hybrid model in the case of replacing one of the backbone outputs with zeros (Fig. 7). Initially, we assumed that the head model would suppress weights of kernel linked with Akita output in case of its uselessness and, therefore, concentrate the weights on VEHiCLE output which shows the good result by itself. However, we observed the opposite behavior: the impact of both backbone outputs was used by the head block to predict the final results. We can speculate that this behavior led all the hybrid models to deteriorate the results.

### 2) Tranformer model for encoder block

We also tried incorporating a sequential Transformer into the sequence-to-image model. Treating DNA data as a ‘language’ seems to be the most natural way. DNA sequences can be represented as a long sequence of chars or split to the short substrings of length k(so-called k-mers). Recently published DNABERT achieved state-of-the-art performance on many sequence prediction tasks. The main difficulty in applying NLP techniques to the DNA analysis is the enormous length of the latter. To some extent, it can be alleviated by Base-Pair Encoding(BPE) as in [CornBERT](https://towardsdatascience.com/bringing-bert-to-the-field-how-to-predict-gene-expression-from-corn-dna-9287af91fcf8). But still, the application of sophisticated Transformer-based architectures is limited because of a lack of resources. We tried to replace the sequence-to-image encoder block constructed from convolutions with the one formed by the DNABERT transformer, but it was impossible to train such a model as there was not enough memory to store all gradients for embeddings computed for the entire input 1Mb sequence. Then, we implemented the model with Akita-like architecture by removing the encoder and feeding the precomputed embeddings from DNABERT to the decoder, but the resulting images became worse while the inference and training time increased. We imply that not only the absence of online augmentation contributed to the poor performance, but the difference between human DNA used for DNABERT training and mouse DNA, which we used as well.

### 3) Data split

Original Akita [21] model was trained on 5 datasets from the human genome with the random split of genomic bins; however, authors of VEHiCLE [20] model used chromosome-based train-validation-test split. We firstly trained our sequence-to-image model on the random split, but then we used the same train-validation-test split for all sequence-to-image, image-to-image, and combined models, as described in Section Data prepocessing. However, the sequence-to-image model shows much better validation scores on random split compared with chromosome-based one (Table III). This fact should be considered for the proper comparison with other methods.

**TABLE III.** Comparison of performances of sequence-to-image model on the random and chromosome-based splits.

| Split | Set | MAE | MSE | Pearson | Spearman |
| --- | --- | --- | --- | --- | --- |
| Random | Train | 0.182 | 0.061 | 0.850 | 0.813 |
|  | Val | 0.246 | 0.115 | 0.682 | 0.656 |
|  | Test | 0.244 | 0.116 | 0.690 | 0.664 |
| Chromosome | Train | 0.220 | 0.092 | 0.778 | 0.760 |
|  | Val | 0.273 | 0.157 | 0.658 | 0.676 |
|  | Test | 0.283 | 0.151 | 0.611 | 0.585 |

### 4) Concluding remarks

Although hybrid models show poor results, we should note that independently trained backbone models were rather successful. The obtained results and learned methodology allow continuing experiments with other types of data to reveal new properties of interconnection between three-dimensional genome form and its nucleotide code.

## Appendix A Code list

- [Pytorch](https://pytorch.org/) – building models.
- [Pytorch-Lightning](https://www.pytorchlightning.ai/) – model training and logging.
- [Torch-geometric](https://github.com/pyg-team/pytorch_geometric) – graph convolution layers.
- [Tensorboard](https://www.tensorflow.org/tensorboard) – logging.
- [Neptune](https://neptune.ai/product) – logging.
- [Akita](https://github.com/calico/basenji/tree/master/manuscripts/akita) – tensorflow code for Akita sequence-to-image model (we adapted it to torch).
- [VEHiCLE](https://github.com/Max-Highsmith/VEHiCLE) – code for VEHiCLE image-to-image model.
- [hicreppy](https://github.com/cmdoret/hicreppy/tree/master/hicreppy) – library for Hi-C specific metric calculation.
- [Cooler](https://github.com/open2c/cooler) – library for Hi-C data reading.
- [cooltools](https://github.com/open2c/cooltools) – library for manipulation of Hi-C data.

## Appendix B Individual contribution

**Dmitrii Kriukov**: overall project design, data preprocessing, literature review, training hybrid models, report writing, presentation preparation, github contributions.

**Mark Zaretckii**: sequence-to-image (Akita) model implementation and training, hybrid model and logging implementation and training, report writing, github contributions.

**Igor Kozlovskii**: sequence-to-image (Akita) model implementation and training, graph models, data preprocessing, report writing, github contribution.

**Mikhail Zybin**: metrics calculation implementation, report writing.

**Nikita Koritskiy**: image-to-image (VEHiCLE) model implementation and training, data preprocessing, comparison with baseline and state of the art models, logging control, report writing, github contributions.

**Mariia Bazarevich**: biological supervising, help with literature review section.

## Appendix C Extra images

## Acknowledgment

The reported study was funded by RFBR, project number 21-34-70051.

